# Corticostriatal stimulation compensates for medial frontal inactivation during interval timing

**DOI:** 10.1101/628263

**Authors:** Eric B. Emmons, Morgan Kennedy, Youngcho Kim, Nandakumar S. Narayanan

## Abstract

Prefrontal dysfunction is a common feature of brain diseases such as schizophrenia and contributes to deficits in executive functions, including working memory, attention, flexibility, inhibitory control, and timing of behaviors. Currently, few interventions can compensate for impaired prefrontal function. Here, we tested whether stimulating the axons of prefrontal neurons in the striatum could compensate for deficits in temporal processing related to prefrontal dysfunction. We used an interval-timing task that requires working memory for temporal rules and attention to the passage of time. Our previous work showed that inactivation of the medial frontal cortex (MFC) impairs interval timing and attenuates ramping activity, a key form of temporal processing in the dorsomedial striatum (DMS). We found that 20-Hz optogenetic stimulation of MFC axon terminals in the DMS shifted response times and improved interval-timing behavior. Furthermore, optogenetic stimulation of terminals modulated time-related ramping of medium spiny neurons in the striatum. These data suggest that corticostriatal stimulation can compensate for deficits caused by MFC inactivation and they imply that frontostriatal projections are sufficient for controlling responses in time.

## Introduction

The prefrontal cortex is dysfunctional in psychiatric disorders such as schizophrenia^1,2^ and neurodegenerative disorders such as Huntington’s disease and Parkinson’s disease^3,4^. Prefrontal impairments are associated with executive dysfunction, including disruption of working memory, attention, flexibility, reasoning, and timing of behavioral responses^5^. Currently, few interventions can mitigate or compensate for prefrontal dysfunction in human disease.

Prefrontal neurons send axons to the basal ganglia^6–9^, and prefrontal impairments can disrupt the function of neurons in subcortical areas^10^. In the present study, we investigated whether manipulating the striatum could compensate for deficits in prefrontal function. To answer this question, we used interval timing, which assesses the ability of subjects to estimate an interval of several seconds based on a motor response. Interval timing is a highly translational function because it involves both prefrontal and striatal regions in rodents as well as in humans and other primates^11–14^ and the neuronal correlates in frontostriatal circuits are similar across species^15–17^. In addition, interval timing is relevant to our research question because it is reliably disrupted in human diseases that impair prefrontal function^16–20^.

A recent study from our group demonstrated that pharmacological inactivation of the rodent medial frontal cortex (MFC; dorsal prelimbic + anterior cingulate)^21,22^ impaired interval timing and attenuated a key neuronal correlate of temporal processing in the dorsomedial striatum (DMS): *time-related ramping activity*^13,23^. These data indicate that temporal processing by DMS neurons requires input via MFC axons, and predict that stimulating MFC➔DMS axon projections can compensate for deficits in temporal control of action as well as striatal temporal processing in animals with the MFC inactivated.

We tested this idea by inactivating the MFC and optogenetically stimulating the terminals of MFC axons in the DMS while rodents performed a 12-second (s) fixed-interval timing task. In animals with intact MFC function, optogenetic stimulation of MFC axon terminals in the DMS had few consistent effects. By contrast, in MFC-inactivated animals, optogenetic stimulation of MFC➔DMS terminals improved response times, curvature of time-response histograms, and time-dependent ramping by DMS neurons. We interpret our results in the context of top-down frontal control of striatal activity, which could be relevant for efforts to mitigate prefrontal dysfunction.

## Results

### Optogenetic stimulation of MFC axons in the DMS normalizes interval-timing behavior

We investigated whether corticostriatal stimulation could compensate for MFC inactivation in 6 rats performing a 12-s interval-timing task. In these rats, both sides of the MFC had been injected with AAV-CamKIIa-ChR2 (MFC-ChR2) and the animals had been implanted with bilateral MFC infusion cannulae and left DMS optrodes. After the rats were acclimatized to infusion and recording procedures, the MFC were bilaterally infused with either saline or the GABAA agonist muscimol, which reversibly and completely inactivates the MFC^13,24,25^. Following this infusion, all animals performed interval timing under three different laser stimulation parameters: no stimulation (No Stim), 2-Hz stimulation (2-Hz), or 20-Hz stimulation (20-Hz). Critically, stimulation was unilateral whereas inactivation was bilateral. We used a generalized linear mixed-effects model (GLMM) to capture effects of optogenetic stimulation on response times in sessions during which the MFC was inactive. Our data reveal a trend for MFC inactivation to affect response times (*F* = 2.9, *p* = 0.09), no effect of optogenetic stimulation, and a significant interaction between MFC inactivation and response time (*F* = 5.1, *p* = 0.006; Table 1; Fig. 2A-B). This interaction was not observed in control animals with AAV-mCherry (interaction: *F* = 0.9, *p* = 0.41).

**Table 1:**
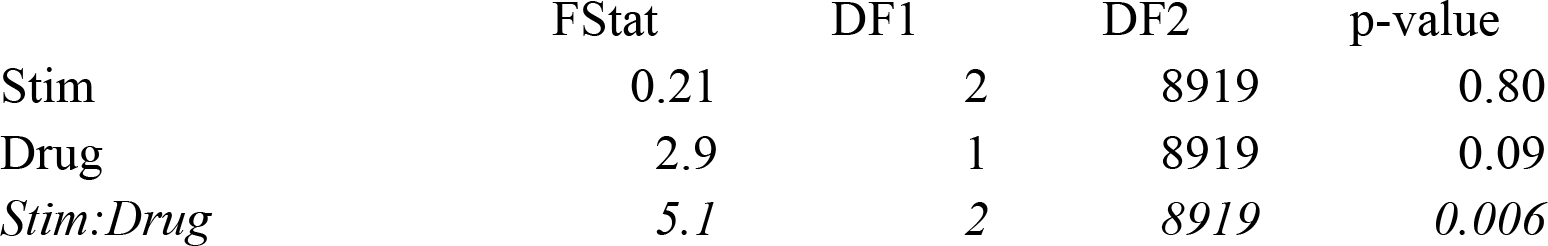
Trial-by-Trial GLME of RT.

**Figure 1.**
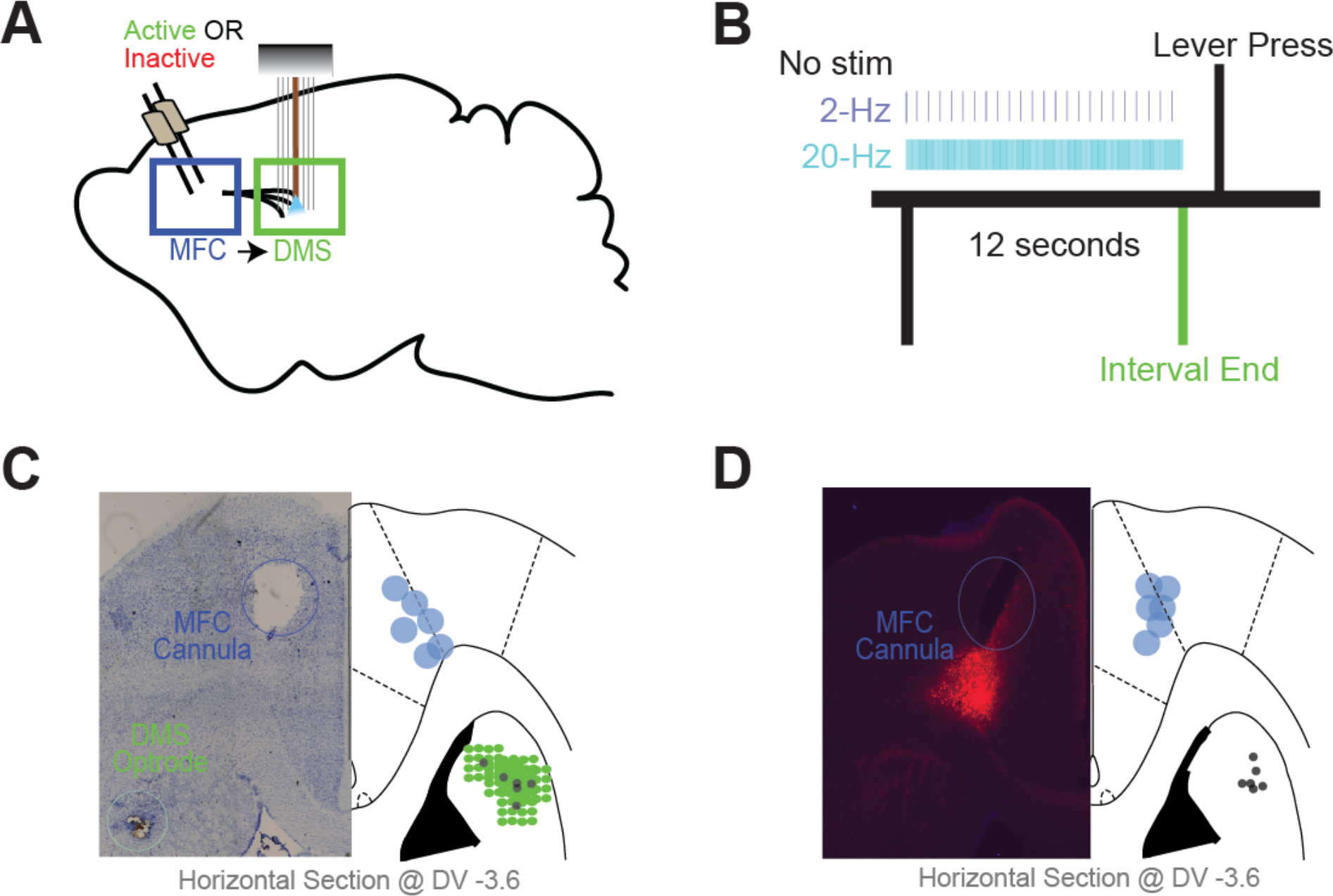
Interval-timing task paradigm and histology. A) Depiction of the surgical approach for the combined optogenetic stimulation and electrophysiological recording experiments. Both MFCs were injected with AAV-ChR2 (or control AAV-mCherry without opsins), after which they were implanted with cannulae along with DMS optrodes (multielectrode recording arrays surrounding a fiber optic cannula). B) Schematic of interval-timing task. A houselight cue signaled the onset of a trial. 12 s later, a reward was available in response to a lever press. MFC➔DMS axons were optogenetically stimulated at 2 Hz or 20 Hz during the interval. C) Left: Representative image of histology in the left hemisphere showing MFC cannula and DMS optrode tracts. Right: Histological reconstruction of placement of cannula and optrode in 6 AAV-ChR2-injected animals. D) Left: Histology of the left hemisphere of a control animal showing MFC cannulae and expression of viral mCherry in the MFC and DMS fiber optics. Right: Histological reconstruction of placement of cannulae and fiber optics in 6 control AAV-mCherry-injected animals.

**Figure 2.**
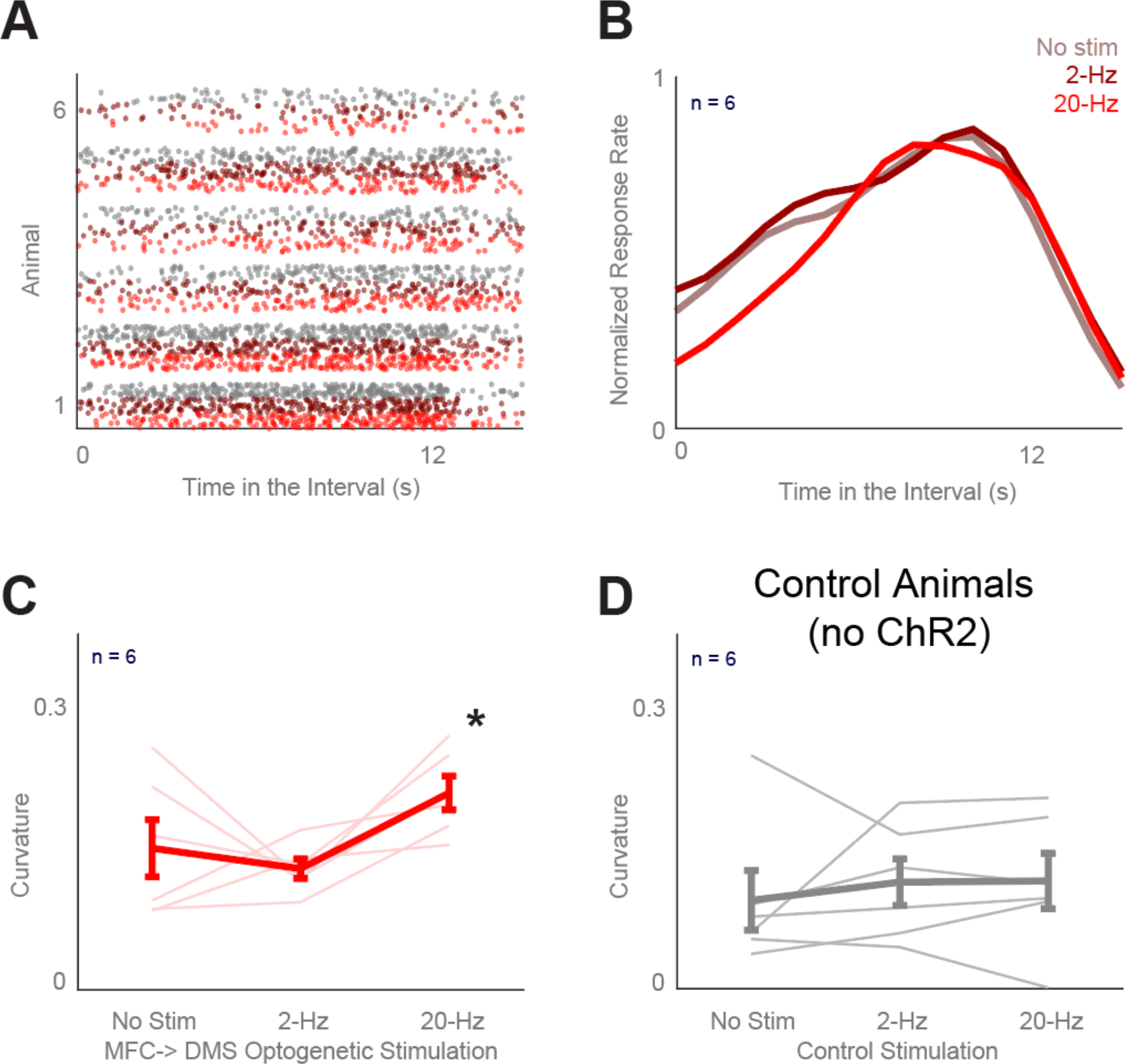
Stimulation of terminals of MFC➔DMS axons influences the timing of responses during interval timing. (A) Raster plots showing all responses of each animal in the study during interval-timing behavior following infusion of the MFC with muscimol. Trials include three laser stimulation conditions: No Stim (grey), 2-Hz (dark red), or 20-Hz (bright red). (B) Average time-response histograms of behavior across laser stimulation conditions (same colors as in A). (C) Average curvature values for animals across stimulation conditions. (D) Average curvature values for control animals (expressing AAV-CamKII-mCherry) across stimulation conditions. All data represent mean ± SEM from sessions in which MFC was inactivated using muscimol; no effects were seen in sessions in which the MFC was infused with saline; \**p* < 0.05.

We also quantified interval-timing performance based on the ‘curvature’ of time-response histograms (Fig. 2C-D)^26,27^. In support of our hypothesis, we found that 20-Hz optogenetic stimulation of the terminals of MFC axons within the DMS increased the curvature of time-response histograms (Fig. 2C; 0.15 ± 0.03 vs. 0.21 ± 0.02; paired *t*_*(5)*_ = 2.6, *p* = 0.05; Cohen’s *d* = 0.95). No significant effects were noted for 2-Hz stimulation or in control animals with AAV-mCherry (Fig. 2C-D). Together, these data provide evidence that optogenetic stimulation of corticostriatal axons at 20 Hz influenced the temporal control of responses primarily when the MFC was inactive.

### Optogenetic stimulation of MFC axons in DMS alters time-dependent ramping in DMS MSNs

Our recent work showed that inactivating the MFC impaired interval timing and attenuated ramping activity in the DMS. Here, we tested the hypothesis that stimulation of MFC axons in the DMS increased time-dependent ramping of DMS neurons. We investigated this question by recording from DMS medium spiny neurons (MSNs; Fig. 3A) while inactivating the MFC and stimulating MFC➔DMS axons. Because the number of fast-spiking interneurons identified by our recording approach was only half that of fast-spiking MSNs, we focused our analyses on MSNs.

**Figure 3.**
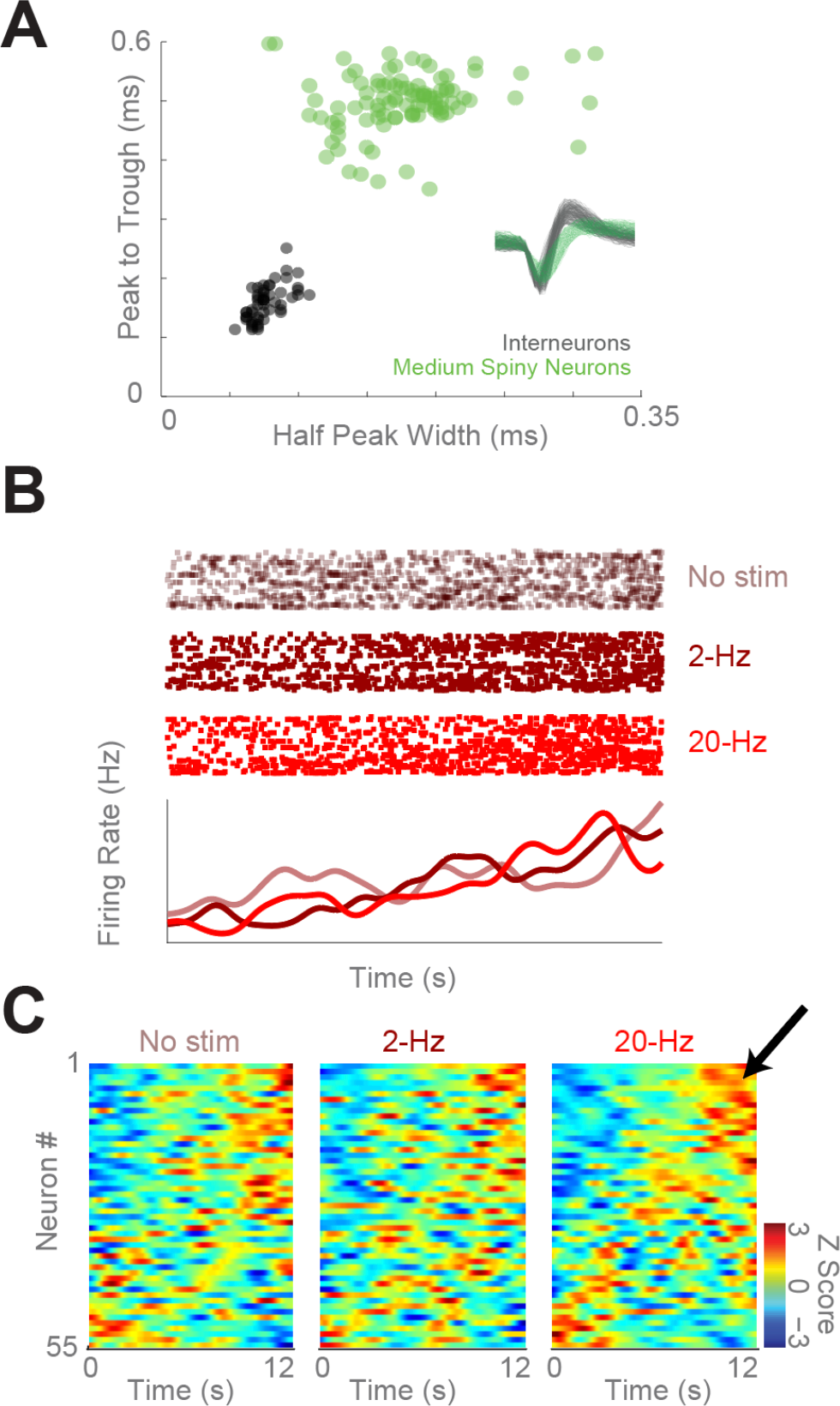
Analysis of MSNs in DMS of animals in which with the MFC is inactive. (A) Separation of medium spiny neurons (MSNs; green) from interneurons (black) in the dorsomedial striatum (DMS) based on peak-to-trough duration and half-peak-width; ms = millisecond. (B) An exemplar DMS MSN with increased time-related ramping with 20-Hz stimulation of MFC➔DMS axons. (C) Peri-event time histograms showing normalized firing rate of all MSNs within the DMS, as recorded during MFC inactivation under each laser stimulation condition. Neurons are sorted based on the first principal component (which, as in our past work, is a ramping pattern of activity). Red indicates higher firing rate than normal, whereas blue indicates lower firing rate. Black arrow highlights increased number of positively-ramping neurons with 20-Hz stimulation.

As in our past work, GLMM analyses revealed that DMS firing rates were affected by optogenetic stimulation (*F* = 8.3, *p* = 0.0003; see Table 2 for stats) and time in the interval (i.e., time-related ramping; *F* = 333.7, *p* = 1e^−74^). Of note, there was an interaction between MFC inactivation and firing rates over time (*F* = 35.6, *p* = 2e^−9^)^13^ and between MFC inactivation and optogenetic stimulation of MFC➔DMS axons (*F* = 5.5, *p* = 0.004, Table 2). Importantly, our model indicated that there was an interaction between time, optogenetic stimulation, and MFC inactivation (*F =* 11.4, *p* = 0.00001; Table 2). These data indicate that optogenetic stimulation changes time-dependent ramping activity as a function of MFC inactivation. This insight was supported by peri-event time histograms which revealed increased time-dependent ramping on trials with 20-Hz optogenetic stimulation of MFC➔DMS axons relative to 2-Hz or No-Stim trials (Fig. 3B-C).

**Table 2:** Trial-by-Trial GLME of MSN firing rate

|  | FStat | DF1 | DF2 | p-value |
| --- | --- | --- | --- | --- |
| *Times* | *333.7* | *1* | *1.6e<sup>7</sup>* | *1.51e<sup>-74</sup>* |
| *Stim* | *8.3* | *2* | *1.6e<sup>7</sup>* | *0.0003* |
| Drug | 0 | 1 | 1.6e<sup>7</sup> | 0.97103 |
| Times:Stim | 1.1 | 2 | 1.6e<sup>7</sup> | 0.34111 |
| *Times:Drug* | *35.6* | *1* | *1.6e<sup>7</sup>* | *2.51e<sup>-09</sup>* |
| *Stim:Drug* | *5.5* | *2* | *1.6e<sup>7</sup>* | *0.004* |
| *Times:Stim:Drug* | *11.4* | *2* | *1.6e<sup>7</sup>* | *0.00001* |

We examined the slope of MSN ramping neurons in MFC-inactivation sessions. The slope was highest on trials with 20-Hz MFC➔DMS axonal stimulation (0.05 ± 0.01 on 20-Hz trials vs. 0.03 ± 0.01 on No-Stim trials, paired *t*_*(50)*_ = 2.5, *p* < 0.01, Cohen’s *d* = 0.26; when restricted to neurons that exhibited ramping: 0.06 ± 0.01 on 20-Hz trials vs. 0.04 ± 0.01 on No-Stim trials; paired *t*_*(38)*_ = 2.1, *p* < 0.05). These analyses provide evidence that the interaction between time-dependent ramping of MSNs, optogenetic stimulation, and MFC inactivation observed in the GLMM analysis was due primarily to increased time-dependent ramping on trials with 20-Hz optogenetic stimulation. Taken together, our results suggest that optogenetic stimulation of MFC axons in the DMS changed both response time and time-dependent ramping activity of DMS neurons as a function of MFC inactivation.

## Discussion

We tested the hypothesis that stimulation of corticostriatal projections could compensate for temporal-processing deficits caused by prefrontal inactivation. We found that in animals in which the MFC was inactivated, stimulation of MFC➔DMS axons at 20 Hz—but not 2 Hz—could normalize the curvature of time-response histograms and shift response times during interval timing. Furthermore, we found that optogenetic stimulation of MFC➔DMS axons interacted with DMS time-dependent ramping activity as a function of MFC inactivation, tending to increase the slope of DMS ramping activity with 20-Hz stimulation in MFC inactivation sessions. Our results provide novel evidence that monosynaptic projections from the MFC to the DMS are sufficient to compensate for behavioral deficits in interval timing caused by MFC inactivation, and also suggest that these projections modulate temporal processing in the DMS.

MFC neurons are involved in top-down control of goal-directed behavior^28–30^. Consistent with our prior work, we found that MFC inactivation decreased the number of MSNs within the DMS that exhibited time-dependent ramping activity, implying that without input from the MFC, a key correlate of temporal processing in the DMS was attenuated. We found that optogenetic stimulation of MFC➔DMS axons could interact with both time-dependent ramping activity and MFC inactivation, providing evidence that stimulation of these axons was sufficient to modulate DMS ramping in MSNs. However, we did not find that optogenetic stimulation increased the number of neurons in the DMS that exhibited time-dependent ramping. Of note, our intervention was relatively restricted in that stimulation was unilateral but pharmacological inactivation was bilateral^31,32^. Whereas MFC inactivation reversibly and completely silences MFC networks^25,33,34^, our optogenetic approach likely stimulated only a small fraction of MFC axons due to the unilateral manipulation and the limitations of viral expression. Furthermore, corticostriatal projections to the striatum are highly organized and specific. Our optogenetic approach nonspecifically stimulated MFC axons. Finally, the activity of MFC neurons during interval timing is highly dynamic^13^, while our stimulation was constant across the interval. In light of these caveats, it is remarkable that the stimulation of corticostriatal terminals was sufficient to influence interval-timing behavior by shifting response times and increasing the curvature of time-response histograms. These data suggest that control of the MFC over DMS ensembles occurs via direct, monosynaptic projections rather than being an indirect effect of MFC projections to another area that modulate the DMS^8,9^.

Here we found stimulation effects at 20 Hz, whereas our past work stimulating MFC networks or MFC afferents revealed effects primarily in the delta/theta range (~4 Hz)^16,17,35,36^. Cortical stimulation can be resonant with ~4-Hz activity among prefrontal networks involved in cognitive control^36^. The striatum may have network properties distinct from cortical rhythms. Importantly, corticostriatal axons release glutamate, and 20-Hz stimulation would release more glutamate at corticostriatal synapses than 2-Hz stimulation^37^. Activity and glutamate release could explain the divergent effects of stimulation frequency on behavior and MSNs. Fast-spiking interneurons can also be modulated by corticostriatal projections, although we recorded fewer such cells and instead focused our analyses on MSNs^38^.

Our study has several additional limitations. Firstly, our approach was limited by the sampling of only a small number of DMS MSNs, and by the nonspecific, virally-mediated expression of opsin in MFC axons that project to the striatum. DMS ramping may depend on specific projections and/or patterns of activity. Additionally, our recording approach did not allow us to differentiate between D1- and D2-dopamine receptor-expressing MSNs or various subtypes of interneurons, and such distinctions might be key to temporal processes^39,40^. Indeed, our recent pharmacological work has suggested that these neuronal populations may perform in distinct ways during interval-timing tasks^41^ and MFC stimulation may preferentially affect one class of neurons vs. the other. Finally, the work presented here was restricted to interval timing, and likely needs to be expanded to other tasks involving sensorimotor and cognitive processing.

In summary, our experiments provide novel evidence that corticostriatal circuits play a critical role in accurate information processing. Combined with our prior work demonstrating that MFC activity is necessary for temporal processing in the DMS, this work suggests that MFC➔DMS projections may be sufficient for some level of rescue of deficits in interval-timing behavior caused by prefrontal dysfunction. We anticipate that this data will inform clinical therapies that target corticostriatal glutamate^42,43^ or deep-brain stimulation of the striatum for disorders that affect the prefrontal cortex^44^.

## Methods

### Rodents

All procedures were approved by the University of Iowa IACUC (protocol #7072039). Twelve male Long-Evans rats were trained on an interval-timing task as described in detail previously^13,25,35,45^. Briefly, animals were initially autoshaped to press a lever for water reward using a fixed-ratio task. Then, animals were trained on a 12-s fixed-interval task (FI12). Trials began with the presentation of a houselight at trial onset (time 0), and the first response made after 12 s had passed resulted in reward delivery, a concurrent click sound, and termination of the houselight (Fig. 1B). Responses made before 12 s elapsed had no programmed consequence. Trials were separated by a randomly chosen intertrial interval of 6, 8, 10 or 12 s. Sessions lasted 60 minutes (m). The timing of each response was used to compute average response rate as a function of time within a trial.

### Surgical and histological procedures

The MFCs of twelve rats were bilaterally infused with AAVs and then implanted with bilateral infusion cannulae, and the left dorsomedial striatum (DMS) was implanted with fiber optic recording electrodes (referred to here as “optrodes”, which were implanted unilaterally). Six of the animals were injected with AAVs that express channelrhodopsin (ChR2); the other six animals were injected with AAVs expressing mCherry and no active channelrhodopsin. Briefly, a surgical level of anesthesia was maintained and, under aseptic surgical conditions, craniotomies were drilled above the left and right MFCs as well as the left DMS. Rat MFCs were first injected with AAV-CamKII-mCherry-ChR2 (AAV-ChR2) or AAV-CamKII-mCherry (AAV-mCherry) virus (1.0 uL virus per side; UNC Viral Vector Core, Chapel Hill, NC). Each rat was later implanted with bilateral infusion cannulae and the left DMS was implanted with a 16-wire optrode (microelectrode array combined with fiber optic cannula). In addition, we used four skull screws. The ground wire from the optrode was connected to the screws and passed into the brain. Optrode arrays consisted of 16 50-μm stainless steel wires arranged in two concentric circles of eight wires surrounding the fiber optic cannula (250 μm between wires and rows; impedance measured *in vitro* at ~400 kΩ; Microprobes for Life Science; Gaithersburg, MD). Infusion cannulae targeted both MFCs (coordinates from bregma: AP +3.2, ML ±1.2, DV −3.6 @ 12° in the anterior plane; these coordinates target the dorsal prelimbic cortex), whereas the optrode recording array targeted only the left DMS (coordinates from bregma: AP +0.0, ML ±4.2, DV −3.6 @ 12° in the lateral plane). Optrode arrays were inserted while recording neuronal activity in order to verify that the implant correctly targeted the DMS. The craniotomy was sealed with cyanoacrylate (‘SloZap’, Pacer Technologies, Rancho Cucamonga, CA), whose polymerization was accelerated by ‘ZipKicker’ (Pacer Technologies); and with methyl methacrylate (i.e., dental cement; AM Systems, Port Angeles, WA). Following implantation, the animals were given one week to recover before being reacclimatized to behavioral and recording procedures.

Following completion of the behavioral experiments, the rats were anesthetized and sacrificed by injection of 100 mg/kg sodium pentobarbital and transcardially perfused with 4% formalin. Brains were post-fixed in a solution of 4% formalin and 20% sucrose before being sectioned on a freezing microtome. Brain slices were mounted on gelatin-coated slides and cell bodies were identified by staining with either DAPI or Cresyl violet. For each animal, histological reconstruction was completed based on postmortem analysis of electrode placements by confocal microscopy or stereology microscopy. These data were used to determine the locations of the electrodes and cannulas within the MFC, and that of the electrode in the DMS (Fig. 1C). Immunohistochemistry was used to visualize the expression of AAV-CamKII-mCherry-ChR2 and AAV-CamKII-mCherry.

### Protocol for rodent behavioral experiments

Rats were first trained in the fixed interval-timing task (FI12) and then underwent stereotactic surgery, as described above. Animals were given one week to recover before being acclimatized to recording and/or stimulation procedures. To ensure that viral expression was maximal, experiments began 3-4 weeks after surgery. Electrophysiological recordings and/or optogenetic stimulation were performed on subsequent days. Infusions of saline and muscimol were performed on separate days. On the first day of the infusion protocol, animals received bilateral saline infusions through both cannulae in the MFC. On the second day, animals were infused with the GABA_A_ receptor agonist muscimol (0.1 mg/mL, 0.5 μL), an approach we have used previously to reversibly and completely inhibit cortical neuronal activity^13,25^. In all recording experiments and/or drug infusions, each session was treated as statistically independent^13,24,25,45–47^. Following the infusions, animal behavior on the FI12 interval-timing task was compared between various conditions of optogenetic stimulation with a 473-nm laser (5-ms pulse width, 10 mW power; Opto Engine, Midvale, UT). Three different stimulation conditions were used in each animal: no stimulation (No Stim), 2-Hz stimulation (2-Hz), and 20-Hz stimulation (20-Hz; **Error! Reference source not found.**A). Conditions were pseudo-randomly interleaved such that the number of trials of each condition (+/− 1 trial) was equal. Behavioral sessions lasted for 60 m.

### Protocol for neurophysiological recordings and analyses of neurons

Neuronal ensemble recordings were made using a multi-electrode recording system (Plexon, Dallas, TX). Putative single neuronal units were identified on-line using an oscilloscope and an audio monitor (Plexon, Dallas, TX). Plexon Offline Sorter was used to analyze the signals after the experiments were completed, and to remove artifacts. Spike activity was analyzed for all cells that fired at rates above 0.1 Hz. Principal component analysis (PCA) and waveform shape were used to sort spikes. Analysis of neuronal activity and quantitative analysis of basic firing properties were carried out using the NeuroExplorer software (Nex Technologies, Littleton, MA) and custom routines available in the MATLAB suite. In each animal, one electrode with minimal neuronal activity was reserved for local referencing, so that 15 electrodes per animal were available for spiking activity. Putative neurons were classified as either medium spiny neurons (MSNs) or fast-spiking interneurons based on waveform peak-to-trough ratio and half-peak widths^48^. MSNs were identified from these parameters by Gaussian mixture clustering in MATLAB (*fitgmdist.m*). Because significantly fewer fast-spiking interneurons were identified, we restricted our analyses to MSNs.

### Statistics

In accordance with our prior work, we quantified temporal control of action by calculating the curvature of time-response histograms. Curvature values range between −1 and 1 and are calculated from the measured cumulative response record by its deviation from a straight line, where 0 would indicate a constant response rate throughout the interval. Curvature indices are resistant to differences in response rate, smoothing, or binning^26,49^.

To quantify the effects of optogenetic stimulation on behavior on a trial-by-trial basis, we used generalized linear mixed-effects models (GLMM; *fitglme* in MATLAB). To quantify the effects of optogenetic stimulation and MFC inactivation on behavior, we used the following GLMM to quantify response time:

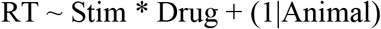

where the dependent variable was response time (*RT*), independent variables were optogenetic stimulation (*Stim:* No Stim, 2-Hz Stim, or 20-Hz Stim) and MFC inactivation (*Drug*), and animals were included as a random variable (*1*|*Animal*). We ran separate models for AAV-ChR2 animals and for AAV-mCherry control animals. For plotting only, kernel density estimates of time-response histograms were calculated. *ksdensity.m* in MATLAB was used with a bandwidth of 1, normalized to the maximum response rate in each animal, and then averaged.

To quantify the effects of optogenetic stimulation, MFC inactivation, and the time of the interval on neuronal activity, we used the following GLMM to quantify neuronal firing rate:

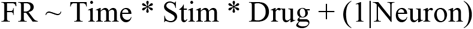

where the dependent variable was firing rate (*FR;* binned at 10 ms over the 12 s interval), and independent variables were time (*Time* over the 12 s interval in 10 ms bins), optogenetic stimulation (*Stim:* No Stim, 2-Hz Stim, or 20-Hz Stim) and MFC inactivation (*Drug*); neurons were included as a random variable (*1*|*Neuron*).

For the analysis of individual neurons, we ran the GLMMs for each neuron of firing rate as a function of time in the interval or of stimulation. This analysis enabled us to identify effects and calculate regression slopes at the single-neuron level. In line with past work, we defined neurons for which the main effect of time had a *p* < 0.05 as exhibiting *time-dependent ramping activity*. All regression analyses were performed on unsmoothed, unnormalized firing rate. For plotting only, peri-event rasters were binned over 10 ms and smoothed over 1 s with a gaussian kernel; peri-event time histograms were binned at 100 ms and smoothed using *ksdensity.m* in MATLAB with a bandwidth of 0.5, normalized using z-scores, and sorted by the first principal component.

## Author contributions

EBE and NSN designed experiments, EBE collected data, EBE, MK, YK, and NSN analyzed data, and EBE, YK, and NSN wrote the paper.

## Data availability statement

All data is available upon request, and a pre-print version of this manuscript will be available on BioRxiv.

## References

1. Andreasen, N. C. et al. Hypofrontality in neuroleptic-naive patients and in patients with chronic schizophrenia. Assessment with xenon 133 single-photon emission computed tomography and the Tower of London. Arch. Gen. Psychiatry 49, 943–958 (1992).

2. Andreasen, N. C. et al. Hypofrontality in schizophrenia: distributed dysfunctional circuits in neuroleptic-naïve patients. Lancet 349, 1730–1734 (1997).

3. Deutch, A. Y. Prefrontal cortical dopamine systems and the elaboration of functional corticostriatal circuits: implications for schizophrenia and Parkinson’s disease. J. Neural Transm. Gen. Sect. 91, 197–221 (1993).

4. Nopoulos, P. et al. Morphology of the cerebral cortex in preclinical Huntington’s disease. Am. J. Psychiatry 164, 1428–1434 (2007).

5. Fuster, J. The Prefrontal Cortex, Fourth Edition. (Academic Press, 2008).

6. Alexander, G. E. & Crutcher, M. D. Functional architecture of basal ganglia circuits: neural substrates of parallel processing. Trends Neurosci. 13, 266–271 (1990).

7. Ferry Amon T., Öngür Dost, An Xinhai & Price Joseph L. Prefrontal cortical projections to the striatum in macaque monkeys: Evidence for an organization related to prefrontal networks. J. Comp. Neurol. 425, 447–470 (2000).

8. Gabbott, P. L. A., Warner, T. A., Jays, P. R. L., Salway, P. & Busby, S. J. Prefrontal cortex in the rat: projections to subcortical autonomic, motor, and limbic centers. J. Comp. Neurol. 492, 145–177 (2005).

9. Han, S.-W., Kim, Y.-C. & Narayanan, N. S. Projection targets of medial frontal D1DR- expressing neurons. Neurosci. Lett. 655, 166–171 (2017).

10. Meyer-Lindenberg, A. et al. Reduced prefrontal activity predicts exaggerated striatal dopaminergic function in schizophrenia. Nat. Neurosci. 5, 267–271 (2002).

11. Matell, M. S. & Meck, W. H. Cortico-striatal circuits and interval timing: coincidence detection of oscillatory processes. Brain Res. Cogn. Brain Res. 21, 139–170 (2004).

12. Coull, J. T., Cheng, R.-K. & Meck, W. H. Neuroanatomical and neurochemical substrates of timing. Neuropsychopharmacol. Off. Publ. Am. Coll. Neuropsychopharmacol. 36, 3–25 (2011).

13. Emmons, E. B. et al. Rodent Medial Frontal Control of Temporal Processing in the Dorsomedial Striatum. J. Neurosci. 37, 8718–8733 (2017).

14. Wang, J., Narain, D., Hosseini, E. A. & Jazayeri, M. Flexible timing by temporal scaling of cortical responses. Nat. Neurosci. 21, 102 (2018).

15. Parker, K. L., Chen, K.-H., Kingyon, J. R., Cavanagh, J. F. & Naryanan, N. S. Medial frontal ~4 Hz activity in humans and rodents is attenuated in PD patients and in rodents with cortical dopamine depletion. J. Neurophysiol. jn.00412.2015 (2015).

16. Parker, K. L. et al. Delta-frequency stimulation of cerebellar projections can compensate for schizophrenia-related medial frontal dysfunction. Mol. Psychiatry 22, 647–655 (2017).

17. Kim, Y.-C. et al. Optogenetic Stimulation of Frontal D1 Neurons Compensates for Impaired Temporal Control of Action in Dopamine-Depleted Mice. Curr. Biol. CB 27, 39–47 (2017).

18. Buhusi, C. V. & Meck, W. H. What makes us tick? Functional and neural mechanisms of interval timing. Nat. Rev. Neurosci. 6, 755–765 (2005).

19. Ward, R. D., Kellendonk, C., Kandel, E. R. & Balsam, P. D. Timing as a window on cognition in schizophrenia. Neuropharmacology (2011).

20. Parker, K. L., Lamichhane, D., Caetano, M. S. & Narayanan, N. S. Executive dysfunction in Parkinson’s disease and timing deficits. Front. Integr. Neurosci. 7, 75 (2013).

21. Laubach, M. A comparative perspective on executive and motivational control by the medial prefrontal cortex. in Neural basis of motivational and cognitive control (MIT Press, 2011).

22. Laubach, M., Amarante, L. M., Swanson, K. & White, S. R. What, If Anything, Is Rodent Prefrontal Cortex? eNeuro 5, (2018).

23. Simen, P., Balci, F., de Souza, L., Cohen, J. D. & Holmes, P. A model of interval timing by neural integration. J. Neurosci. Off. J. Soc. Neurosci. 31, 9238–9253 (2011).

24. Narayanan, N. S., Horst, N. K. & Laubach, M. Reversible inactivations of rat medial prefrontal cortex impair the ability to wait for a stimulus. Neuroscience 139, 865–876 (2006).

25. Parker, K. L., Chen, K.-H., Kingyon, J. R., Cavanagh, J. F. & Narayanan, N. S. D1-Dependent 4 Hz Oscillations and Ramping Activity in Rodent Medial Frontal Cortex during Interval Timing. J. Neurosci. 34, 16774–16783 (2014).

26. Fry, W., Kelleher, R. T. & Cook, L. A mathematical index of performance on fixed-interval schedules of reinforcement. J. Exp. Anal. Behav. 3, 193–199 (1960).

27. Narayanan, N. S., Land, B. B., Solder, J. E., Deisseroth, K. & DiLeone, R. J. Prefrontal D1 dopamine signaling is required for temporal control. Proc. Natl. Acad. Sci. U. S. A. 109, 20726–20731 (2012).

28. Niki, H. & Watanabe, M. Cingulate unit activity and delayed response. Brain Res 110, 381–6 (1976).

29. Ostlund, S. B. & Balleine, B. W. Lesions of medial prefrontal cortex disrupt the acquisition but not the expression of goal-directed learning. J Neurosci 25, 7763–70 (2005).

30. Ma, L., Hyman, J. M., Phillips, A. G. & Seamans, J. K. Tracking progress toward a goal in corticostriatal ensembles. J. Neurosci. Off. J. Soc. Neurosci. 34, 2244–2253 (2014).

31. Xu, R. et al. Quantitative comparison of expression with adeno-associated virus (AAV-2) brain-specific gene cassettes. Gene Ther. 8, 1323–1332 (2001).

32. Hommel, J. D., Sears, R. M., Georgescu, D., Simmons, D. L. & DiLeone, R. J. Local gene knockdown in the brain using viral-mediated RNA interference. Nat. Med. 9, 1539–1544 (2003).

33. Martin, J. H. & Ghez, C. Pharmacological inactivation in the analysis of the central control of movement. J Neurosci Methods 86, 145–59 (1999).

34. Allen, T. A. et al. Imaging the spread of reversible brain inactivations using fluorescent muscimol. J. Neurosci. Methods 171, 30–38 (2008).

35. Emmons, E. B., Ruggiero, R. N., Kelley, R. M., Parker, K. L. & Narayanan, N. S. Corticostriatal Field Potentials Are Modulated at Delta and Theta Frequencies during Interval-Timing Task in Rodents. Front. Psychol. 7, (2016).

36. Kim, Y.-C. & Narayanan, N. S. Prefrontal D1 Dopamine-Receptor Neurons and Delta Resonance in Interval Timing. Cereb. Cortex N. Y. N 1991 (2018).

37. Kreitzer, A. C. Physiology and Pharmacology of Striatal Neurons. Annu. Rev. Neurosci. 32, 127–147 (2009).

38. Burguiere, E., Monteiro, P., Feng, G. & Graybiel, A. M. Optogenetic Stimulation of Lateral Orbitofronto-Striatal Pathway Suppresses Compulsive Behaviors. Science 340, (2013).

39. Tecuapetla, F., Matias, S., Dugue, G. P., Mainen, Z. F. & Costa, R. M. Balanced activity in basal ganglia projection pathways is critical for contraversive movements. Nat. Commun. 5, (2014).

40. Howard, C. D., Li, H., Geddes, C. E. & Jin, X. Dynamic Nigrostriatal Dopamine Biases Action Selection. Neuron 93, 1436–1450 (2017).

41. De Corte, B. J., Wagner, L. M., Matell, M. S. & Narayanan, N. S. Striatal dopamine and the temporal control of behavior. Behav. Brain Res. 356, 375–379 (2019).

42. Emre, M. et al. Memantine for patients with Parkinson’s disease dementia or dementia with Lewy bodies: a randomised, double-blind, placebo-controlled trial. Lancet Neurol. 9, 969–977 (2010).

43. Garcia-Munoz, M., Lopez-Huerta, V. G., Carrillo-Reid, L. & Arbuthnott, G. W. Extrasynaptic glutamate NMDA receptors: key players in striatal function. Neuropharmacology 89, 54–63 (2015).

44. Dougherty, D. D. et al. A Randomized Sham-Controlled Trial of Deep Brain Stimulation of the Ventral Capsule/Ventral Striatum for Chronic Treatment-Resistant Depression. Biol. Psychiatry 78, 240–248 (2015).

45. Parker, K. L., Ruggiero, R. N. & Narayanan, N. S. Infusion of D1 Dopamine Receptor Agonist into Medial Frontal Cortex Disrupts Neural Correlates of Interval Timing. Front. Behav. Neurosci. 9, 294 (2015).

46. Narayanan, N. S., Cavanagh, J. F., Frank, M. J. & Laubach, M. Common medial frontal mechanisms of adaptive control in humans and rodents. Nat. Neurosci. 16, 1888–1897 (2013).

47. Narayanan, N. S. & Laubach, M. Neuronal correlates of post-error slowing in the rat dorsomedial prefrontal cortex. J. Neurophysiol. 100, 520–525 (2008).

48. Berke, J. D., Okatan, M., Skurski, J. & Eichenbaum, H. B. Oscillatory Entrainment of Striatal Neurons in Freely Moving Rats. Neuron 43, 883–896 (2004).

49. Narayanan, N. S., Land, B. B., Solder, J. E., Deisseroth, K. & DiLeone, R. J. Prefrontal D1 dopamine signaling is required for temporal control. Proc. Natl. Acad. Sci. 109, 20726–20731 (2012).

